# A function of Spalt proteins in heterochromatin organization and maintenance of genomic DNA integrity

**DOI:** 10.1101/2024.04.30.591908

**Authors:** Cristina M. Ostalé, Natalia Azpiazu, Ana Peropadre, Mercedes Martín, Mireya Ruiz-Losada, Ana López-Varea, Rebecca R. Viales, Charles Girardot, Eileen E.M. Furlong, Jose F. de Celis

## Abstract

The phylogenetically conserved Spalt proteins regulate gene expression and participate in a variety of cell fate choices during multicellular development, generally acting as transcriptional repressors in different gene regulatory networks. Paradoxically, besides their roles as DNA sequence-specific transcription factors, Spalt proteins show a consistent localization to heterochromatic regions. They can act through interactions with the Nucleosome remodeling and deacetylase complex (NuRD) to promote closing of open chromatin domains, but their activities as epigenetic regulators also rely on interactions with DNA Methyltransferases or with the Lysine-specific histone demethylase LSD1, suggesting that they can participate in multiple regulatory mechanisms. Here we describe several major consequences of loss of *spalt* function in *Drosophila* cells, including changes in chromatin accessibility affecting mostly pericentromeric heterochromatin, the generation of DNA damage, alterations in the localization of chromosomes within the nucleus in polyploid cells of the salivary glands and miss-expression of transposable elements. We suggest that most of these effects are related to roles of Spalt proteins in the regulation of heterochromatin formation. We propose that *Drosophila* Spalt proteins have two complementary functions, acting as sequence-specific transcriptional repressors on specific target genes and regulating more global gene silencing through the generation or maintenance of heterochromatic domains.

## Introduction

The Spalt proteins (Sal) constitute a conserved family of nuclear proteins whose function have been related to the regulation of gene expression, acting as sequence-specific transcription factors (de Celis and Barrio 2009; Sweetman and Münsterberg 2006). They are characterized by the presence of between 7 and 10 C2H2 Zinc finger domains and a poli-Q stretch, and contain several sites of Sumoylation related with Sal activity and subnuclear localization (Sánchez et al. 2010, 2011). The Sal proteins play multiple seemingly unrelated roles during the development of vertebrates and invertebrates, including vulva formation and mesodermal and neural cell fate specification in *C. elegans* (Grant et al. 2000; Jarriault et al. 2008; Shen et al. 2017; Toker et al. 2003), and limb, heart, kidney and neuronal development in vertebrates (Álvarez et al. 2021; Sweetman and Münsterberg 2006). The *Drosophila sal* genes, *spalt-major* (*salm*) *and spalt-related* (*salr*), form part of a gene complex (Barrio et al. 1999), and like other *sal* genes their functions are required for multiple developmental processes including the acquisition of segmental identity, the development of the Johnston organ, tracheal dorsal trunk formation, the specification of some photoreceptor cells in the eye, mesoderm and neuronal development, and imaginal wing disc growth and patterning (de Celis et al., 1996; Domingos et al., 2004; Eberl and Boekhoff-Falk, 2007; Kühnlein and Schuh, 1996; Kuhnlein et al., 1994).

The diversity of roles played by Sal proteins in processes of cell fate allocation is in part due to their participation in different transcriptional gene regulatory networks operating in different tissues and developmental times (Martín et al. 2016). Within these networks, the Sal proteins have a broad impact in the transcriptional profile of cells, altering, directly or indirectly, the expression of a multitude of genes in mutant conditions. This has been shown during the development of the murine kidney (Basta et al., 2014), odontoblasts (Lin et al. 2021), embryonic stem cells (Miller et al. 2016)(El Pantier et al. 2021), the *Drosophila* wing disc (Organista et al., 2015) and in processes of tumoral transformation (Yang et al. 2017). The multiplicity of functional requirements for Sal proteins in the assignation of cell fates, and the broad changes in gene expression observed in different mutant conditions for the *sal* genes, underly the association of human SALL1 and SALL4 with the genetic syndromes Townes-Brocks (SALL1; Kohlhase 2000; Kohlhase et al. 1999) and Okihiro (SALL4; Borozdin et al. 2004), which are characterized by a variety of malformations in the limbs and different internal organs. Similarly, the roles of SALL proteins in the maintenance of pluripotency (Yang et al. 2008; Sakaki-Yumoto et al. 2006a; Tsubooka et al. 2009), may explain the implications of these genes with a variety of human cancers (Álvarez et al. 2021).

The mechanisms by which the Sal proteins regulate gene expression have been the subject of considerable research interest, but there is still not a unifying framework to understand their functions as transcriptional regulators. On the one hand, Sal proteins directly regulate cell-type specific target genes by binding to regulatory sequences. For example, *Drosophila* Salm/Salr directly represses *knirps* (*kni*) acting through putative enhancers located 11 Kb from the *kni* transcription start site (Ostalé et al. 2024) and the *C. elegans* Sal-like protein Sem-4 represses the expression of the *egl-5* and *mec-3* genes during neuronal differentiation, and this regulation requires the binding of Sem-4 to regulatory sequences located in the promotors of its target genes (Toker et al. 2003). Similarly, mouse SALL2 regulates cell proliferation in mouse embryonic fibroblast by acting as a direct repressor of *cyclin D1* and *cyclin E1* (Hermosilla et al., 2018). Further examples of sequence-specific transcription factor activity have been reported for SALL4 on *HOXA9* expression during human hematopoiesis (Gao et al. 2013), on glycolytic enzymes in murine limbs (Kawakami et al. 2023), and for the repression by Sall1/Sall4 of *pou5f3* in the case of *Xenopus* neural patterning (Exner et al. 2017). Taken together, these results indicate that SAL proteins have a conserved role as direct transcriptional repressors, binding in a sequence specific manner to repress enhancers or promoters.

On the other hand, Sal proteins preferentially locate to heterochromatic regions and interact with several heterochromatic proteins, suggesting an activity of Sal proteins related to the organization of the heterochromatin and potentially to the regulation of gene expression by epigenetic modifications (Netzer et al. 2001, 2006; Sakaki-Yumoto et al. 2006). In addition, numerous studies delineate the pivotal involvement of the Nucleosome Remodeling and Deacetylase (NuRD) complex as a central facilitator in the repression mediated by SALL1 and SALL4 (Kiefer et al. 2002; Wang et al. 2023; Lauberth et al. 2007; Lauberth and Rauchman 2006). This mechanism of gene regulation does not operate in all cell contexts requiring Sal function, as the activity of NuRD is not required for Sall1 and Sall4 transcriptional repression of neuronal genes in mouse embryonic stem cells (Miller et al. 2016). Finally, Sall4’s role as a transcriptional repressor has also been linked to its recruitment of the lysine-specific histone demethylase LSD1, which mediates the conversion of di-methylated H3K4 to mono- and unmethylated H3K4 (Liu et al. 2013). The variety of Sal interaction partners and mechanisms of transcriptional regulation in which these proteins participate suggest a model in which Sal proteins recruit epigenetic complexes to either cell-type specific gene target promotors or to particular chromatin domains to regulate gene expression or chromatin structure.

Here we analyzed the possible contributions of *Drosophila* Sal proteins to chromatin organization as a mechanism to account for the variety of *sal* developmental roles during wing imaginal development. We find that loss of *salm* and *salr* expression results in considerable changes in chromatin accessibility in imaginal cells, but that these changes bear little relation to the changes in gene expression detected in the same genetic background. In addition, these changes mostly affect pericentromeric heterochromatic regions, which are the regions preferentially bound by endogenous Salm proteins in imaginal cells (Ostalé et al. 2024). We also found profound changes in the nuclear localization of chromosomes within the nucleus in polyploid cells of the salivary glands, and an unexpected relationship between Sal function and the maintenance of genome integrity and transposable elements silencing. We suggest that these effects are due to loss of heterochromatin formation and/or maintenance in *sal* mutant cells. Altogether, our results draw a scenario in which diploid imaginal *sal* mutant cells display altered heterochromatin assembly and/or organization, suffer mitotic errors and DNA damage, trigger DNA-damage responses and fail to progress to mitosis, whereas *sal* mutant polyploid cells have defects in nuclear envelope morphology and nucleolar size accompanied by changes in the nuclear disposition of chromatin. We suggest that these functions of Sal proteins underly the majority of changes in gene expression observed in *sal* mutant cells, and define a global role for these proteins in heterochromatin maintenance that complements their activities as a sequence-specific transcription factor acting on a specific set of target genes.

## Results

### Loss of spalt function has a profound impact on chromatin accessibility

The analysis of global mRNA expression in *salm/salr* mutant wing discs revealed a multitude of changes in the expression levels of a large set of genes, both increases and decreases in gene expression (Organista et al., 2015). Here, we analyzed variations in chromatin accessibility in wild type discs compared to *salm/salr* knockdown discs, aiming to find possible correlations between changes in gene expression levels and chromatin accessibility in these two genetic backgrounds. To define chromatin accessibility, we carried out ATAC-seq (“Assay for Transposase-Accessible Chromatin with sequencing”) experiments in control discs (*sal^EPv^-Gal4/UAS-GFP*) and in *salm/salr* knock-down discs (*sal^EPv^-Gal4/UAS-salm-RNAi; UAS-salr-RNAi/+*). At an FDR (False Discovery Rate) corrected p-value of 0.05, we detected 5175 chromatin regions with accessibility changes (Fig. 1A and Supple. Table 1). The number of regions that increase or reduce their accessibility conformation in *salm/salr* knock-down conditions compared to wild type discs is similar (2760 vs 2416, respectively; Fig. 1A; Supple. Table 1). At a more stringent corrected p-value FDR (0.001), the number of regions with significant changes in DNA accessibility is still high, and includes 899 and 642 regions that increase or decrease their accessibility, respectively (Fig. 1A).

**Figure 1:**
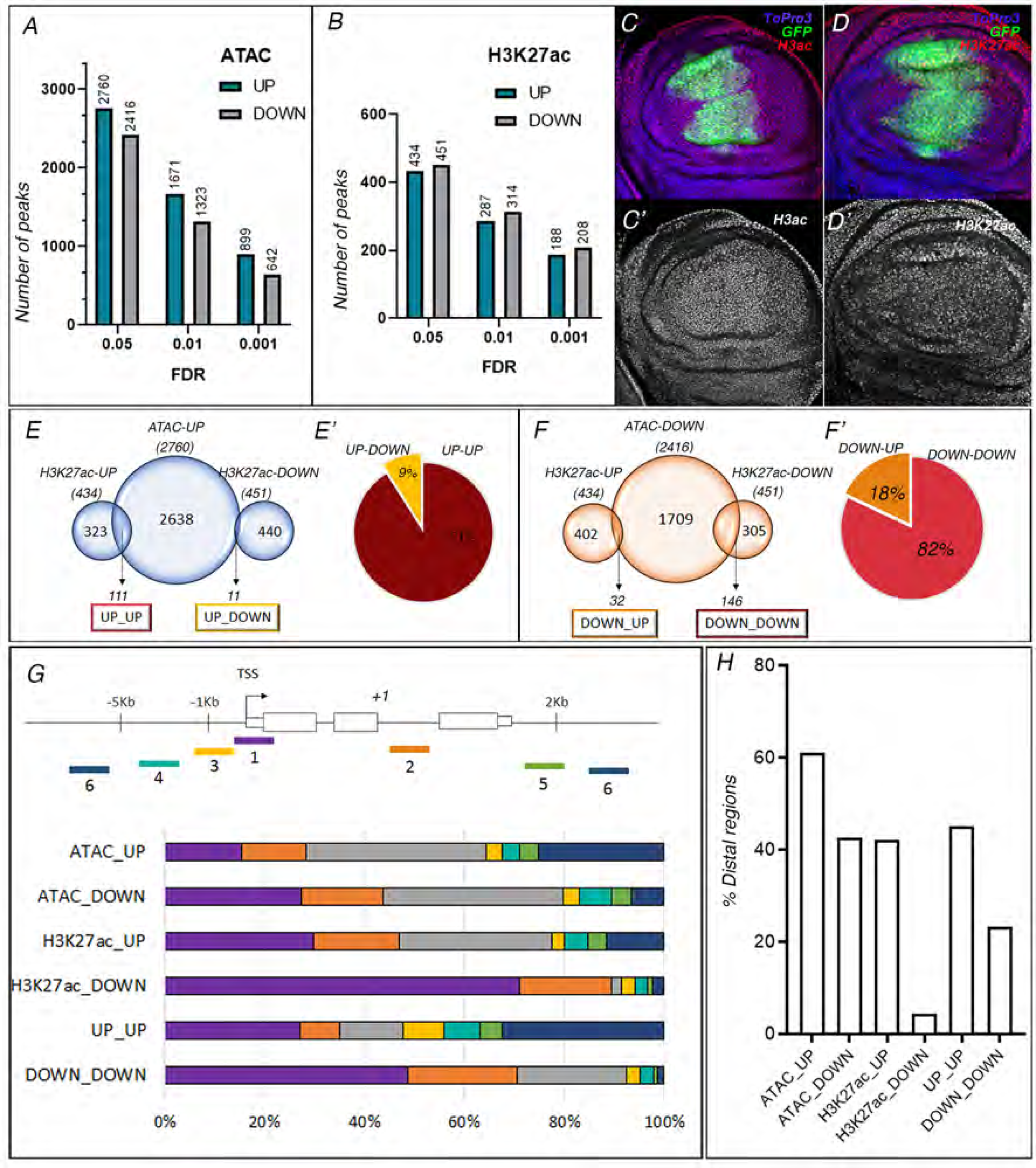
Changes in chromatin accessibility in *salm/salr* knockdown imaginal discs. (A-B) Number of genomic regions showing a significant enrichment (UP, green) or depletion (DOWN; grey) comparing the ATAC-seq data sets (A) or H3K27ac ChIP-seq data sets (B) of *salm/salr* mutant wing imaginal discs (*sal^EPv^-Gal4/UAS-salm-RNAi; UAS-salr-RNAi/+*) and control wing discs (*sal^EPv^-Gal4/UAS-GFP*). The DNA regions were identified at 0.05, 0.01 and 0.001 false discovery rates (FDR). (C-D) Expression of H3ac (C) and H3K27ac (D) in late third instar wing imaginal discs of *sal^EPv^-Gal4/UAS-salm-RNAi; UAS-salr-RNAi/UAS-GFP* genotype. The expression of GFP is shown in green, the expression of H3ac (C) and H3K27ac (D) in red and the expression of DAPI in blue. The individual red channels of C and D are shown in C’ and D’, respectively. (E-E’) Comparison between the genomic regions showing a significant enrichment in the ATAC-seq data set (ATAC-UP; FDR 0.05) with those enriched or depleted in the H3K27ac ChIP-seq data set (H3K27ac-UP and H3K27ac-DOWN, respectively). The percentage of H3K27ac-UP and H3K27ac-DOWN regions included in the ATAC-UP class is shown in E’. (F-F’) Comparison between the genomic regions showing a significant depletion in the ATAC-seq data set (ATAC-DOWN; FDR 0.05) with those enriched or depleted in the H3K27ac ChIP-seq data set (H3K27ac-UP and H3K27ac-DOWN, respectively). The percentage of H3K27ac-UP and H3K27ac-DOWN regions included in the ATAC-DOWN class is shown in F’. (G) Proximity criteria to classify genomic regions (above; 1: promoter, 2: proximal intragenic, 3: proximal intergenic, 4: medial intergenic, 5: proximal 3’, 6: distal intergenic; in grey are represented regions located within large introns) and percentage of each sequence class in the different data sets. (H) Percentage of intragenic and intergenic distal sequences present in each data set.

Active DNA regulatory regions are enriched for certain Histone post-transcriptional modifications such as H3 acetylation in lysine 27 (H3K27ac; Bonn et al. 2012; Fu et al. 2018). We found that loss of *salm/salr* function also has a broad impact on the presence of the H3K27ac mark in imaginal cells. Thus, ChIP-seq (“Chromatin immunoprecipitation followed by sequencing”) experiments with anti-H3K27ac antibodies in control and *salm/salr* knock-down discs reveal that, at a corrected p-value FDR 0.05, there are 434 and 451 chromatin regions showing increase or decrease, respectively, acetylation in H3K27 (Fig. 1B; Supple. Table 1). As expected from these numbers, the overall accumulation of H3K27ac in wing discs of *sal^EPv^-Gal4/UAS-salm-RNAi; UAS-salr-RNAi/+* genotype is not affected (Fig. 1C-D). The overlap between changes in accessibility and changes in H3K27ac, such as ATAC-UP with H3K27ac-UP or H3K27ac-DOWN (Fig. 1E), and ATAC-DOWN with H3K27ac-UP or H3K27ac-DOWN (Fig. 1F), is low. For example, only 122 of the ChIP-seq H3K27ac regions (14%) are included in the ATAC-UP class (Fig. 1E). However, 91% of these 122 regions that gain accessibility (ATAC-UP) correspond to increases in H3K27 acetylation (Fig. 1E). Similarly, only 178 regions with modifications in H3K27ac (20%) are included in the ATAC-DOWN class (Fig. 1F). In this case, most regions of overlap (82%) correspond to those showing reduced H3K27 acetylation (Fig. 1F).

When we associated the sequences found in the ATAC-seq and H3K27ac ChIP-seq with their nearest gene affected (Fig. 1G; see material and methods), we found a large number of enriched sequences located more than 5 Kb from the nearest transcription start site (TSS), including those located within large introns (distal intragenic) or between genes (distal and proximal intergenic; Fig. 1G). The fraction of these distal regions is maximal in the ATAC-UP class (61%), and this fraction is below 42% only for those regions included in the H3K27ac-DOWN and H3K27ac-DOWN/ATAC-DOWN classes (Fig. 1H). A detailed analysis of the genomic location of the DNA regions altered in the ATAC-seq and H3K27ac ChIP-seq sequences revealed a prominent localization close to the centromeres (Fig. 2A-C). The pericentromeric regions are mostly formed by heterochromatin, large stretches of highly repetitive and densely packed DNA with low gene density, which includes approximately 30% of the *Drosophila* genome (180 Mb; He et al., 2012). We analyzed the pericentromeric heterochromatin regions as defined by the epigenetic frontiers identified by the accumulation of the H3K9me2 and H3K9me3 marks (Riddle et al. 2011) and by cytology in the case of the chromosomal arm 3R (Hoskins et al. 2007). Including the autosomes and the X chromosome, this pericentromeric heterochromatin includes 14.1% of the genome (around 19Mb out of 134Mb). Analyzing the sets of regions that suffer epigenetic modification after *salm/salr* knockdown, we found a particular enrichment in pericentromeric areas within the regions that gain accessibility in *salm/salr* knockdown discs (22.1% of total bp whose accessibility is increased with a corrected p-value FDR 0.001; Fig. 2A-C). These results suggest a role for Sal proteins in pericentromeric chromatin organization, specifically in the maintenance of chromatin with low accessibility.

**Figure 2:**
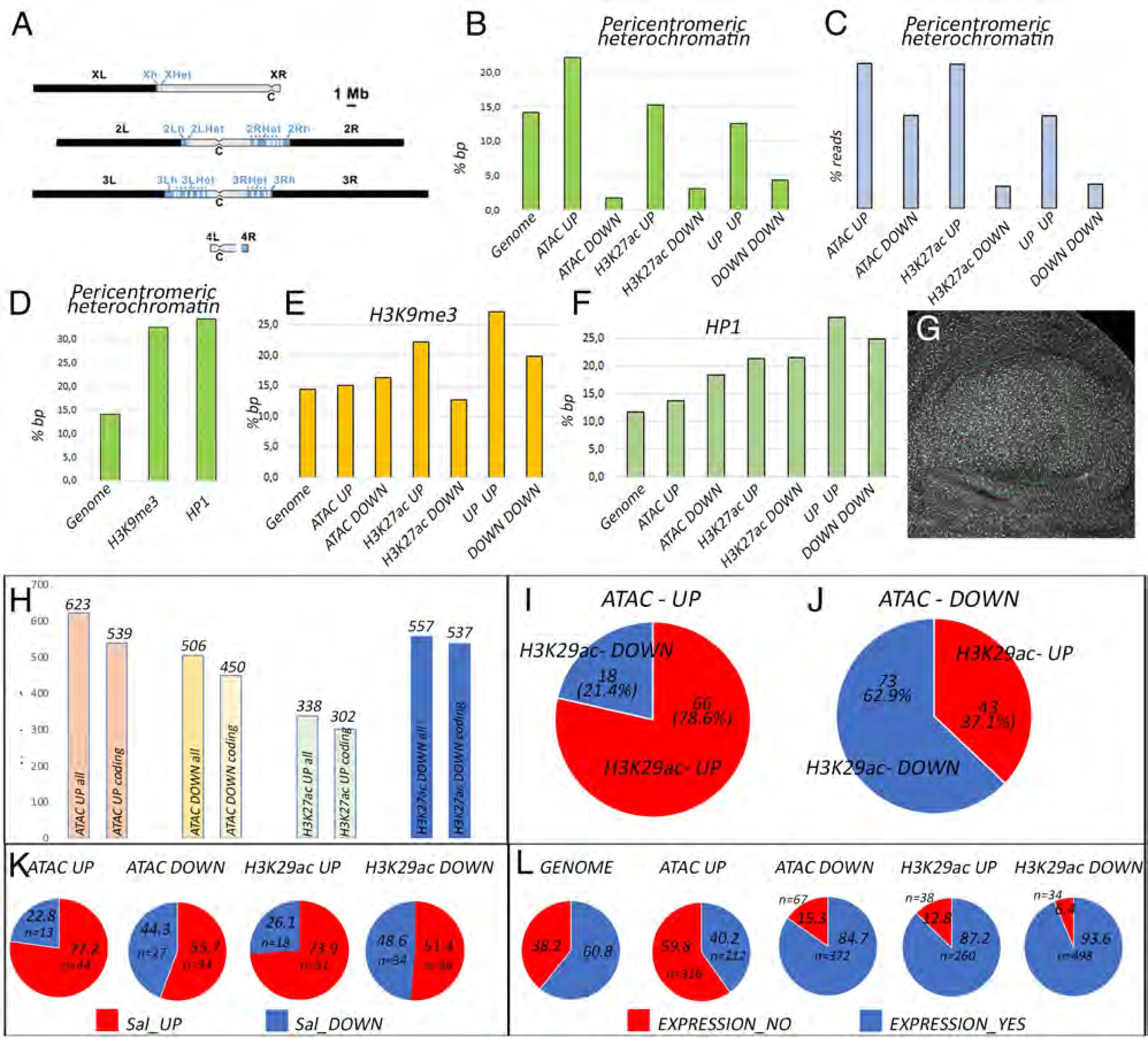
Association of chromatic accessibility and H3K27as modifications changes observed in *salm/salr* mutant discs with heterochromatin regions. (A) Distribution of pericentromeric heterochromatin (blue regions in the grey centromeric domain, C) and euchromatin (black) in *Drosophila* chromosomes. (B-C) Percentages of sequence lengths (B) and sequence reads (C) located in pericentromeric chromatin for the genome (Genome; B) and for all the sequences identified in ATACseq (ATAC UP and ATAC down) or H3K27ac ChIP seq (H3K27ac UP and H3K27ac DOWN) experiments. (D) Percentage of sequence lengths (% bp) in the genome and detected in H3K9me3 and HP1 ChIP seq datasets. (E-F) Percentage of sequence length identified in ATACseq and H3K27ac ChIP seq that are also associated to H3K9me3 (E) and HP1 (F) ChIP seq datasets. (G) Expression of H3K9me3 in a wing imaginal disc of *sal^EPv^-Gal4 UAS-GFP/UAS-salm-RNAi; UAS-salr-RNAi* genotype. The area of *sal^EPv^-Gal4* expression is delineated with a green line. (H) Number of genes (all) and coding genes (coding) associated to the sequences identified in ATACseq and H3K27ac ChIP seq experiments. (I-J) Percentage of genes included in the data sets ATAC-UP (I) and ATAC-DOWN (J) included in the datasets H3K29ac-DOWN (blue sectors) or H3K29ac-UP (red rectors). (K) Percentage of overlap between genes which expression is augmented (Sal_UP; red) or reduced (Sal_DOWN; blue) in *salm/salr* mutant discs included in the ATAC UP, ATAC DOWN, H3K29ac UP and H3K29ac DOWN datasets. (L) Percentage of genes normally expressed in the wild type wing disc (EXPRESSION_YES; blue) or not expressed in the wild type wing disc (EXPRESSION_NO (red) in the entire genome (GENOME) or in the ATAC UP, ATAC DOWN, H3K29ac UP and H3K29ac DOWN datasets.

Constitutive heterochromatin, present in telomeric and centromeric regions of the chromosomes, is characterized by the presence of epigenetic repression marks such as the trimethylation of H3K9 or the association of Su(var)205 (HP1), which are involved in the establishment and maintenance of the conformational state of this chromatin form (Swenson et al., 2016). More than 30% of the sequence coverage of H3K9me3 and HP1 ChIP-sequencing (ChIP-Seq H3K9me3: Oregon L3, Karpen, G. Dataset ID: 4952 and ChIP-Seq HP1: Oregon L3, Karpen, G. Dataset ID: 4936) align to pericentromeric regions (Fig. 2D). Because of this shared genomic location enrichment, we explored whether the regions that suffer chromatin activity and accessibility changes after *salm/salr* knockdown are usually bound by HP1 and/or are located in H3K9me3 modified landscapes. We found a consistent overlap between HP1-bound fragments and all regions whose open state or H3K27ac abundance is modified after *salm* and *salr* RNAi expression. For all sets (ATAC_UP, ATAC_DOWN, H3K27ac_UP and H3K27ac_DOWN), the proportion of nucleotides sequenced in each set that is also sequenced after HP1 chromatin immunoprecipitation is significantly higher than the 11.8 % of bp bound by HP1 in the genome (values of 15 % to 22 %; Fig. 2F). Regarding H3K9me3 bound sequences, we observed a specific enrichment in those regions that increased its accessibility and activation state (H3K27ac abundance) after *salm/salr* silencing (Fig. 2E). The occurrence of epigenetic changes after *salm/salr* knockdown in wing discs, especially in regions where HP1 is bound in wildtype conditions, suggests a possible function of Sal proteins in the regulation or maintenance of chromatin architecture.

We also compared the sets of genes most likely affected by these epigenetic modifications (see Fig. 2H). There is only a weak overlap between the associated genes with changes in H3K27ac and ATAC peaks (Fig. 2I-J and Supple. Table 1). Thus, 200 out of the 846 genes associated to changes in H3K27ac are included within the 997 genes associated to changes in accessibility (Fig. 2I-J). Within these 200 genes, there is a much better correspondence between genes with increased H3K27ac and higher chromatin accessibility (66/84 genes; 78%), and genes with lower H3K27ac and lower accessibility (73/116 genes; 63%; Fig. 1I-J).

The candidate genes associated with changes in H3K27ac and/or changes in ATAC peaks are not highly correlated with the genes that changes in expression in *salm/salr* knockdown(Organista et al. 2015). To do this comparison, we used a set of 1622 genes whose expression changes significantly (pvalue lower than 0.05) after 24-or 48-hour periods of *salm/salr* knockdown, irrespectively of their fold change (Supple. Table 1). We found that only 257 of these genes were also present in either the differential H3K27ac or ATAC associated genes (Fig. 2K). Considering all possible pairwise comparisons between the three datasets (Fig. 2K) we noticed that the best overlaps were those including genes upregulated in *salm/salr* knock-down discs acquiring higher accessibility and enriched in H3K27ac labeling (Fig. 2K). Genes associated to regions of higher chromatic accessibility in *salm/salr* knock-down discs generally correspond to genes that are not expressed in the wing discs in wild-type conditions (Fig. 2L). On the contrary, genes associated to loss of accessibility or to changes in H3K27ac modifications are expressed in imaginal discs at a much higher proportion than observed for the entire genome (Fig. 2L). In conclusion, we find that *salm/salr* knockdown wing discs display numerous changes in chromatin accessibility and histone modifications that are enriched in distal regions to the transcription start sites and pericentromeric heterochromatin regions. Heterochromatic regions are also preferentially bound by Salm in Salm ChIP-seq experiments (Ostalé et al. 2024). However, a large fraction of changes in ATACseq and H3K27Ac are not correlated with the changes in gene expression observed in *salm/salr* knockdown wing imaginal discs.

### Requirements of salm/salr in the maintenance of heterochromatic regions

The pattern of Salm binding to pericentromeric DNA (Ostalé et al., 2024), as well as the alterations in chromatin conformation affecting pericentromeric chromatin (Fig. 2A-C), suggest a role for Sal proteins in the generation and/or maintenance of heterochromatic regions. Mutations in genes encoding proteins required for heterochromatin formation are often haplo-insufficient, and in heterozygous conditions increase the expression of genes placed into heterochromatic positions (Elgin and Reuter 2013; Swenson et al. 2016). To assess if this is also the case for *sal*, we first checked whether epigenetic silencing occurs in *salm/salr* heterozygous cells by looking at the expression of *miniwhite* insertions present in the subtelomeric regions of the 2R and 3R chromosomic arms. In both cases we found a consistent increase in eye pigmentation comparing P{hsp26-pt-T} w[67c23] expression in females heterozygous for a deficiency for the *salm* and *salr* genes (*Df(2L)32FP-5*) with their corresponding *CyO* siblings (Fig. 3A-E). We also looked at retrotransposon expression of the HeT-A, TART and F-element families in 0-2 hours old embryos. The F-element is mainly localized in the heterochromatin of the fourth chromosome, whereas the HeT-A and TART retrotransposons are the main components of *Drosophila* telomers (Casacuberta and Pardue 2005;Riddle et al. 2008). In all three cases we found an average four-fold increase in the expression of the retrotransposons in the progeny of *Df(2L)32FP-5* heterozygous flies compared with wild type (*y w*) embryos (Fig. 3F). We also looked at the nuclear localization of the heterochromatin associated protein HP1 (Kellum et al. 1995). HP1 mostly localizes in the polyploid nuclei of salivary glands in a single spot that corresponds to the chromocenter, and is also present in several discrete chromosomal arms and the telomeres (Fig. 3G). The localization of HP1 in the nuclei of *salm/salr* knockdown salivary glands appears in several spots located in proximity to the nucleolus (Fig. 3H). Taken together, these results suggests that cells heterozygous for the *salm/salr* deficiency have defective silencing in the expression of retrotransposons and white^+^ insertions in telomeric regions, as well as alterations in the pattern of HP1 accumulation within the nucleus.

**Figure 3:**
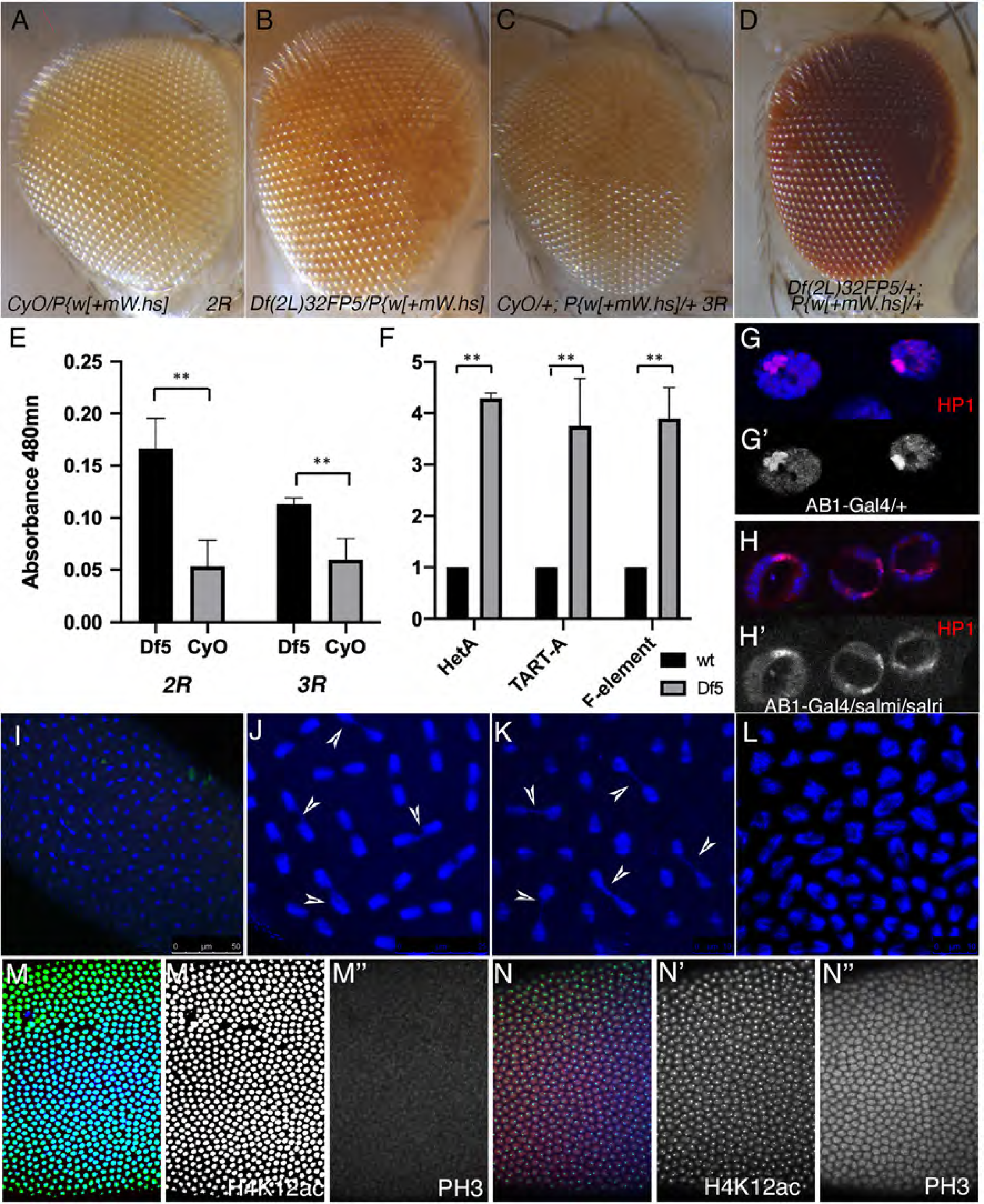
Changes in gene expression in heterochromatic regions in *salm/salr* heterozygous flies and embryos. (A-B) Two days old eyes prepared from males of *CyO/P{w[+mW.hs]* (A) and *Df(2L)32FP5/P{w[+mW.hs]* (B) genotypes. The insertion P{w[+mW.hs] is located in the Telomere Associated Sequence of the 2R chromosomic arm. (C-D) Two days old eyes prepared from males of *CyO/+; P{w[+mW.hs]*/+ (C) and *Df(2L)32FP5/+; P{w[+mW.hs]/+* (D) genotypes. The insertion P{w[+mW.hs] is located in the Telomere Associated Sequence of the 3R chromosomic arm. (E) Eye pigment quantification (measured in absorbance at 480 Mn) of *Df(2L)32FP5/P{w[+mW.hs]* (Df5; black) and *CyO/P{w[+mW.hs]* (CyO; grey) in the first two left columns, and *Df(2L)32FP5/+; P{w[+mW.hs]/+* (Df5; black) and *CyO/+; P{w[+mW.hs]*/+ (CyO; grey) in the two right columns. (F) Expression (qPCR) of the retrotransposons HetA, TART-A and F-element in control embryos (black columns) and in embryos obtained from *Df(2L)32FP5* heterozygous flies. (G-H) Distribution of HP1 (red) and DAPI (blue) in salivary gland nuclei from larvae of *AB1-Gal4 UAS-GFP/+* (G-G’) and *AB1-Gal UAS-GFP/UAS-salm-RNAi; UAS-salr-RNAi/+* (H-H’). The corresponding red channels are shown in G’ and H’. (I) Confocal images of mitotic chromosomes (DAPI, blue) of a syncytial blastoderm from heterozygous *Df(2L)32FP5* parents. (J-L) Higher magnification of independent visual fields showing defective mitotic figures (anaphase bridges, white arrowheads in J-K) and asynchronic mitotic figures (L). (M-M’’) Distribution of H4K12ac (green in M and white in M’), Phospho-Histone3 (PH3, red in M and white in M’’), and DAPI (blue in M) in wild type blastoderm (cycle 12). (N-N’’) Expression of H4K12ac (green in N and white in N’), Phospho-Histone3 (PH3, red in N and white in N’’), and DAPI (blue in N) in a the embryonic blastoderm (cycle 12) derived from heterozygous *Df(2L)32FP5* flies.

The correct formation of heterochromatin in pericentromeric and telomeric regions is a prerequisite for the correct segregation of chromosomes during mitosis. We monitored mitotic chromosomes in early blastoderm embryos (0-2 h AEL) obtained from the progeny of flies heterozygous for a deficiency including the *salm* and *salr* genes (*Df(32FP-5*; Barrio et al. 1999). At this stage, mitosis of blastoderm nuclei are synchronic in normal embryos (Foe 1989), but we found a variety of alterations in the embryos arising from *Df(32FP-5* heterozygous flies (Fig. 3I-L). These alterations include the formation of anaphase bridges (Fig. 3J-K), loss of synchrony in the mitotic figures and a general disorganization of the mitotic spindles (Fig. 3L). We also observed significant changes in the expression of Histone modifications (H4K12ac and phospho-Histone3) in 0-2 h old embryos from *Df(2L)32FP-5/CyO* mothers (Fig. 4M-N). The H4K12ac modification, which interferes with PH3 deposition (Boros 2012), is expressed at very low levels in *salm/salr* mutant blastoderms, presenting an abnormal distribution within the nucleus (Fig. 3M-M’, N-N’). As a consequence, PH3 is detected at higher-than-normal levels in all *salm/salr* mutant preblastodermal nuclei in interphase, before they start mitosis (Fig. 3M-M’’ and N-N’’). All these data indicate that in early blastoderms the reduction in Salm/Salr protein levels is associated with changes in chromatin organization, histone modifications and the appearance of mitotic errors.

**Figure 4:**
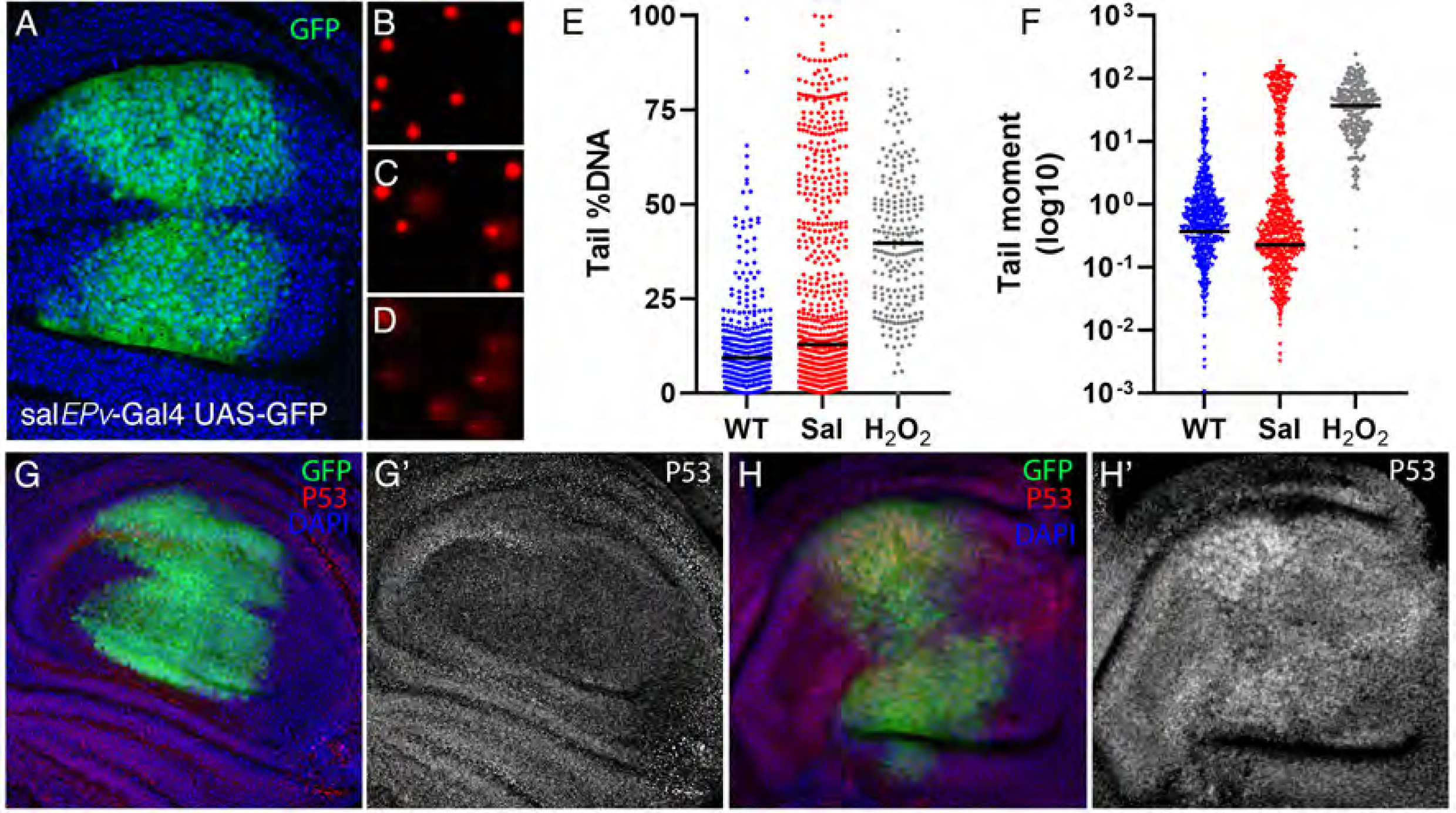
DNA damage in *salm/salr* mutant wing imaginal discs. (A) Expression of GFP (green) in the wing blade region of *sal^EPv^-Gal4 UAS-GFP/+* late third instar larva. The expression of DAPI is shown in blue. (B-D) Representative examples of wing imaginal disc nuclei obtained from larvae of *sal^EPv^-Gal4 UAS-GFP/+* (B), *sal^EPv^-Gal UAS-GFP/UAS-salm-RNAi; UAS-salr-RNAi/+* (C) and *sal^EPv^-Gal4 UAS-GFP/+* dissociated discs treated for 10 min with H_2_O_2_ 50 µM (D). (E-F) Percentage of Tail DNA (left) and value of Tail moment (right) in the single cell electrophoresis assay from *sal^EPv^-Gal4 UAS-GFP/+* (blue dots), *sal^EPv^-Gal UAS-GFP/UAS-salm-RNAi; UAS-salr-RNAi/+* (red dots) and *sal^EPv^-Gal4 UAS-GFP/+* exposed to H_2_O_2_ (grey dots). (G-H) Expression of P53 (red in G and H) in control discs (*sal^EPv^-Gal4 UAS-GFP/+*; G-G’) and in *sal^EPv^-Gal UAS-GFP/UAS-salm-RNAi; UAS-salr-RNAi/+* discs (H-H’). The expression of GFP is in green (G-H) and DAPI in blue (G-H). The individual red channels (P53) are shown in G’-H’.

### Salm/Salr function is required for genome integrity

Loss of heterochromatin silencing can also lead to DNA damage and to the activation of DNA replication check points, altering the progression of the cell cycle (Fortuny and Polo 2018). Interestingly, *salm/salr* mutant wing discs show a prominent accumulation of cells in the G2 phase of the cell cycle and a loss of mitotic cells (Organista and de Celis, 2013), which is compatible with the existence of DNA damage. In order to determine the existence of DNA damage in imaginal cells deficient for the *salm/salr* genes, we made use of the single cell electrophoresis assay for detecting single and double-strand DNA breaks in dissociated wing imaginal discs of *sal^EPv^-Gal4/UAS-salm-RNAi; UAS-salr-RNAi/+* genotype. The expression of *sal^EPv^-Gal4* is restricted to the central domain of the wing blade (Fig. 4A). Consequently, cells dissociated from these discs contains *salm/salr* knockdown cells (wing cells from the domain of *salm/salr* expression), normal wing cells lacking expression of the *salm/salr* genes, and cells from the wing hinge and thorax that express *salm/salr* but do not express the corresponding RNAi’s, and therefore have normal Salm and Salr function. In these experiments we used wild type discs (*sal^EPv^-Gal4/UAS-GFP)* as a negative control, to provide information about background physiological levels of DNA damage. As a positive control we used wing discs extracted from *sal^EPv^-Gal4/UAS-GFP* larvae treated with H_2_O_2_ 50 µM for 10 min, as the exposure of cells to peroxide very effectively induces DNA double and single-strand breaks (Driessens et al. 2009). Fluorescent imaging of DNA from single nucleus readily showed the existence of a cell population of cells displaying DNA damage that is present in *salm/salr* knockdown discs but not in control *sal^EPv^-Gal4/UAS-GFP* discs (Fig. 4B-C). In contrast, wing discs treated with H_2_O_2_ showed a unique population of cells with considerable DNA damage (Fig. 4D). As expected from these images, the quantification of the percentage of DNA in the tails (Fig. 4E), as well as the values of Tail moment (Fig. 4F), a measure that incorporates the amount and the extent of DNA damage in cells, confirm the existence of DNA damage in *salm/salr* mutant discs (Fig. 4E-F).

A key player in the coordination of DNA damage responses is the tumor suppressor protein P53, whose protein levels increase as a response to DNA damage (Hafner et al. 2019). Once activated by phosphorylation, p53 regulates the expression of a multitude of genes related to cell division, cell death and DNA repair (Hafner et al. 2019). The expression of P53 is detected at similar levels in all cells of the wildtype wing disc (Fig. 4G-G’). In contrast, we found that the levels of P53 protein are increased in the central region of the wing blade in *salm/salr* mutant discs (Fig. 4H-H’), further confirming the existence of DNA damage cells in the domain of Salm/Salr expression. Taken together, our results suggest that an important consequence of the loss of *salm/salr* function in the wing disc is the generation of DNA damage, which could be linked to the defects in cell cycle progression observed in *salm/salr* mutant cells, and to a role of these genes in the generation or maintenance of heterochomatin.

### Loss of salm/salr compromises nuclear envelope morphology and results in increased nucleolar size in salivary gland polyploid cells

Heterochromatic regions appear mostly associated with the inner nuclear envelop and the periphery of the nucleolus (Bizhanova and Kaufman 2021). In fact, heterochromatin packing contributes to the maintenance of the stiffness of the nuclear envelope, nucleolus morphology and the normal function of the nuclear pore complexes (Ballmer et al. 2023; Shevelyov 2023; Bustin and Misteli 2016). To analyze in more detail the possible effects of *salm/salr* functions on chromatin, we studied the consequences of *salm/salr* knockdowns in the salivary gland, a secretory epithelium formed by large polyploid cells (Denisa et al. 2021; Haberman et al. 2003). The *spalt* genes are expressed in the nuclei of all salivary gland cells (Fig. 5A), and loss of *salm/salr* function in these cells causes a variety of phenotypes, ranging from rudimentary salivary glands to morphologically normal glands, depending on the particular Gal4/UAS-RNAi combination (Fig. 5B-E). We focused our analysis in the salivary glands of *salm/salr* hippomorphic conditions (weak knockdown) because they have an overall morphology similar to wildtype glands. Salm/salr salivary glands have smaller cells with nuclear sizes smaller than those of normal glands (Fig. 5B-E). Interestingly, the nuclear envelope is undulated, and presents numerous indentations, in contrast with normal salivary gland nuclei, which present a straight nuclear envelope without major indentations (Fig. 5F-G). In addition, the size of the nucleolus is much larger in *salm/salr* mutants compared to normal glands, and the chromosomes have an abnormal disposition within the nuclei (Fig. 5H-I). These defects are very similar to those found in prothoracic glands mutant for *salm*/*salr* (Ostalé et al. submitted), suggesting a general requirement of Salm/Salr function, at least in polyploid cells, to maintain the normal disposition of heterochromatin within the nucleus and to regulate, most likely indirectly via the changes in heterochromatin, morphological aspects of the nuclear envelope and nucleolus. These requirements could correspond to what has been defined as non-genetic aspects of the genome, through which its physical and mechanical properties affect morphological and functional aspects of the nucleus (Bustin and Misteli 2016).

**Figure 5:**
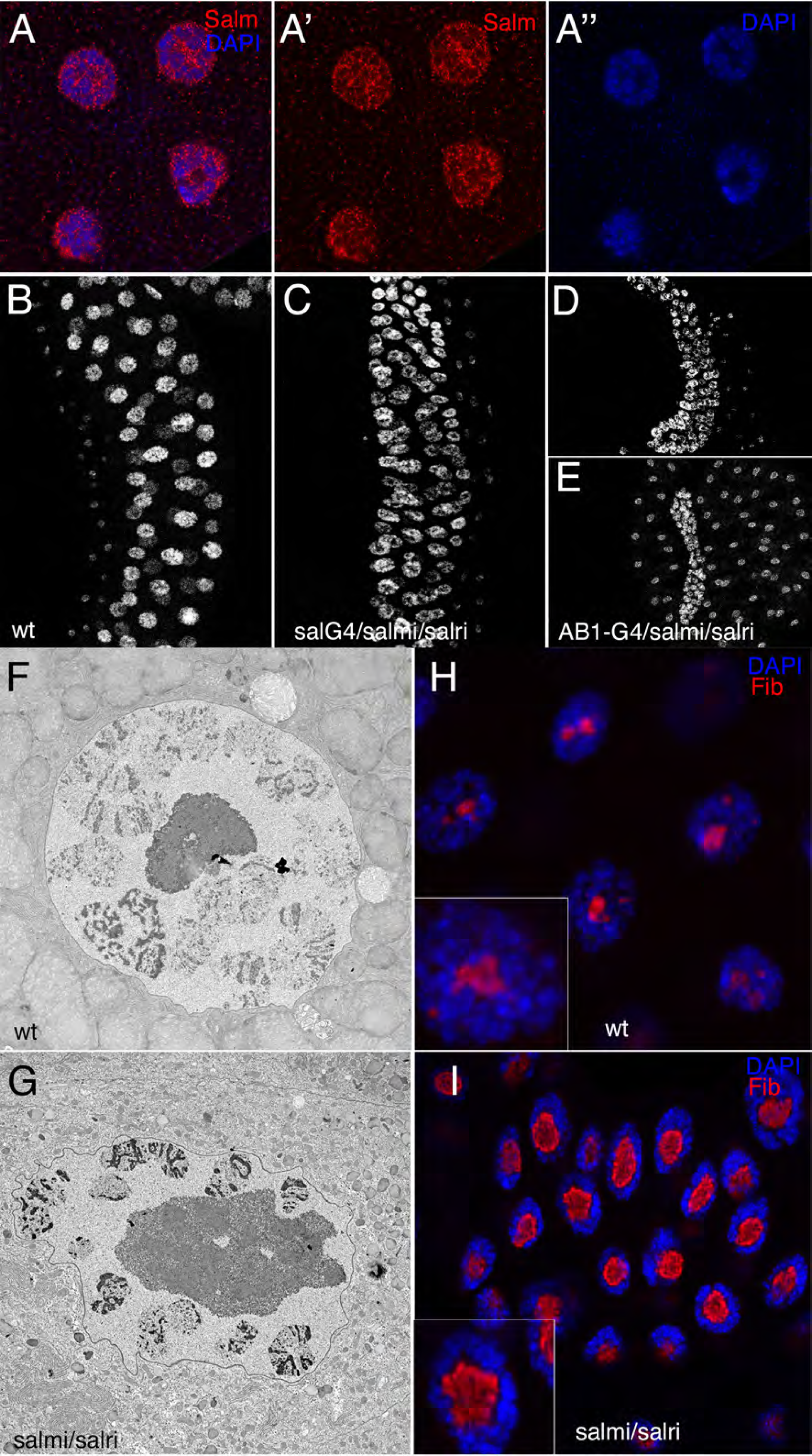
Nuclear morphology in the salivary gland of *salm/salr* knockdown larvae. (A-A’’) Expression of Salm (red in A-A’) and DAPI (blue in A and A’’) in wild type salivary glands from late third instar larvae. (B-E) Wild type and *salm/salr* mutant salivary glands of different genotypes. Each panel shows the expression of DAPI in salivary glands from control larvae (*sal^EPv^-Gal4 UAS-GFP/+*; B), *sal^EPv^-Gal4 UAS-GFP/UAS-salm-RNAi; UAS-salr-RNAi/+*; C) and *UAS-salm-RNAi/+; AB1-Gal4/UAS-salr-RNAi* (D-E). (F-G) TEM images of the cell nucleus in a wild type salivary gland (F) and in a *sal^EPv^-Gal4 UAS-GFP/UAS-salm-RNAi; UAS-salr-RNAi/+* salivary gland. Note the different size of the nucleolus and the alterations in the shape of the nuclear membrane. (H-I) Control (H) and *sal^EPv^-Gal4 UAS-GFP/UAS-salm-RNAi; UAS-salr-RNAi/+* salivary glands showing the expression of Fibrillarin (Fib; red) and DAPI (blue). Insets in each panel are magnifications (3x) showing single nucleus.

### Irradiation and loss of Salm/Salr cause overlapping changes in gene expression in the wing disc

Our results above suggest that a prominent function of the Salm/Salr proteins is related to the establishment and/or maintenance of heterochromatic regions within the nucleus, and that a major consequence of Salm/Salr loss is the generation of DNA damage. To assess this, we searched for links between the transcriptional changes observed in *salm/salr* mutant discs (FDR 0.05, logFC < −0.4 and >0.4; (Organista et al., 2015) and those occurring as a consequence of DNA damage induced by irradiation (van Bergeijk et al. 2012). We noticed a reasonably overlap between the two datasets, particularly for those genes which expression increases after irradiation and in *salm/salr* mutant discs (Fig. 6A and Supple Table 2). We also searched for changes in expression levels after irradiation of a set of genes that became overexpressed in the central region of *salm/salr* mutant discs genes (Organista et al., 2015). Most of these genes (86%; 25/29) correspond to genes overexpressed after irradiation with logFC in the range of 1 to 6 (Fig. 6B-C). Interestingly, a considerable number of these genes (9/25) encode proteins involved in DNA repair (Fig. 6B). In two cases where we identified the enhancer region of the corresponding gene that became active upon a loss of *salm/salr* function (*Gadd45* and *ver;* Ostalé et al., 2024), the same enhancers also became activated after irradiation (Fig. 6D-G). The increase in expression levels observed after irradiation depends on the transcriptional regulator Dp53 (van Bergeijk et al. 2012), which is expressed in all wing cells at low levels even upon irradiation (Fig. 6H-I).

**Figure 6:**
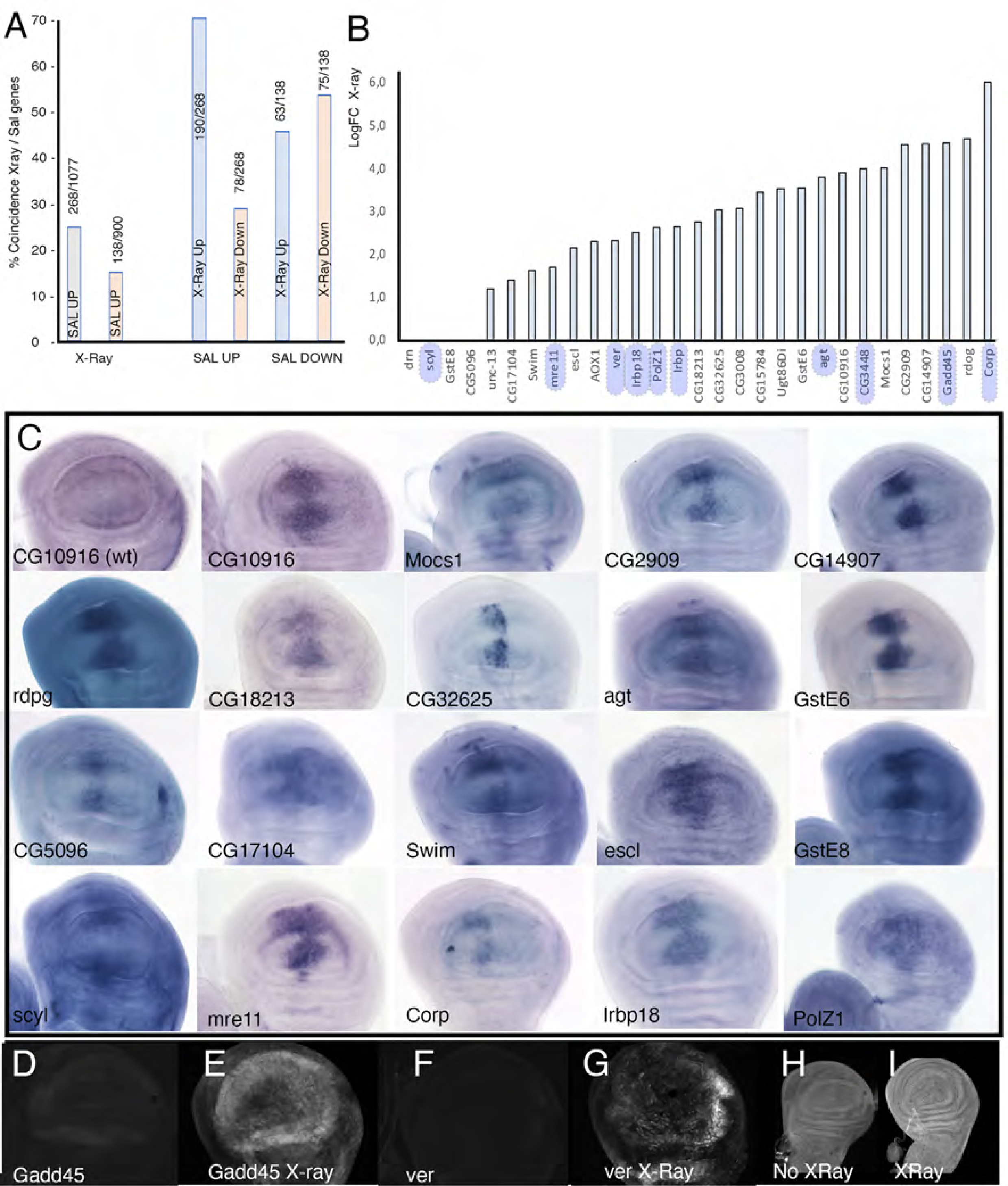
Gene expression analysis in *salm/salr* knockdown and irradiated wing imaginal discs. (A) Two left columns: Percentage of genes which expression changes in *salm/salr* mutant discs and in irradiated wing discs (van Bergeijk et al. 2012). Four right columns: Percentage of genes which expression is increased (SAL UP) or reduced (SAL DOWN) that were also identified as showing increased (X-Ray Up) or reduced (X-ray Down) expression after irradiation of wild type wing imaginal discs. Each number indicates the number of genes in each class. (B) LogFC values of changes in gene expression after irradiation of wild type wing imaginal discs (van Bergeijk et al. 2012) for a collection of genes identified in *salm/salr* mutant discs showing ectopic expression in the central domain of the wing disc. All genes within blue boxes are related to DNA damage responses. (C) Late third instar wing imaginal discs showing the mRNA expression by *in situ* hybridization of the indicated gene (bottom left corner of each panel) in a *sal^EPv^-Gal4/UAS-salm-RNAi; UAS-salr-RNAi* genetic background. Only the expression of CG10916 in wild type wing discs is shown in the first ‘panel. All other genes are either not expressed or expressed in all cells at low levels in wild type wing discs. (D-E) Expression of the regulatory region of *Gadd45* (genomic coordinates 2R:7245799-7249134 (+)) in third instar wing imaginal discs of *Gadd45 RR-GFP/+* genotype. The larvae were grown in normal conditions (D) or irradiated with 3000 R 2 h before dissection (E). Expression of the regulatory region of *ver* (genomic coordinates 3L:12523886-12524405(+)) in third instar wing imaginal discs of *ver RR-GFP/+* genotype. The larvae were grown in normal conditions (F) or irradiated with 3000 R 2 hours before dissection (G). (H-I) Expression of P53 in wild type wing discs (H) and in wild type wing discs irradiated with 3000 R 2 hours before dissection (I).

## Discussion

We have identified a variety of cellular and genetic alterations in cells mutant for the *Drosophila salm* and *salr* genes that suggest a novel aspect of their function as transcriptional regulators. We focused mostly on the wing imaginal disc, a tissue where the developmental requirements of these genes have been extensively characterized. During imaginal wing disc development, *salm/salr* are required for the specification of the thorax, the patterning of veins and sensory organs, and also to promote cell viability, epithelial integrity and the cell cycle transition from G2 to mitosis (Organista and de Celis, 2013; Grieder et al., 2009, Martín et al., 2017; Ostalé et al., 2024). It is assumed that these phenotypes are somehow caused by miss-regulation of Sal target genes, but only some aspects of *sal* function have been linked to the direct regulation of specific target genes. For example, the function of Salm/Salr during the positioning of the presumptive longitudinal L2 vein relies on a gene regulatory network including the transcriptional repression of *knirps* and *Optix* by Salm/Salr (Martín et al. 2017), and at least *knirps* repression involves binding of Salm to a regulatory sequence located 11 Kb from the *knirps* transcription start site (Ostalé et al. 2024). Similarly, the function of Salm/Salr in the regulation of epithelial integrity could be mediated by repression of *tartan* and *capricious* expression in the central region of the developing wing blade (Milán et al. 2002). In contrast, no candidate genes have been identified mediating the functions of Salm/Salr required to promote cell cycle progression and cell viability in the wing blade. Complementary, loss of *salm/salr* function generates a major change in the transcriptional landscape of the wing disc (Organista et al., 2015), but the relation between these changes and the known functional requirements of Salm/Salr is not apparent. It is also unknown to what extent the observed changes in gene expression characteristic of *salm/salr* mutant discs are a consequence of Salm/Salr acting as canonical transcription factors on multitude of target genes.

### Loss of salm/salr function alters DNA accessibility in wing disc cells

A new insight into the transcriptional effects of *salm/salr* loss of function conditions came from the analysis of chromatin accessibility and the characteristics of the Salm binding pattern to the DNA. We found that the knockdown of *salm* and *salr* genes in wing imaginal discs display massive changes in chromatin accessibility, which affect DNA regions preferentially located more than 5 Kb distal from transcription start sites or included in pericentromeric heterochromatin domains. We also detected changes in the presence of the H3K27ac histone mark in *salm* and *salr* knockdown wing imaginal discs, but these were less numerous that alterations in chromatin accessibility, suggesting that remodeling of chromatin in enhancer sequences could be a less determining event in the transcriptional response to Salm/Salr. The overlap between the two data sets was only enriched for regions that simultaneously increased in both chromatin accessibility and H3K27ac modifications, as well as for those showing the opposite change, less accessibility and less acetylation of H3K27. Consistently with these data, we recently found that Salm binding to the DNA is enriched in pericentric heterochromatin, as approximately 80% of chromatin bound by Salm belongs to this class (Ostalé et al. 2024). All together, these observations indicate that a prominent function of Salm/Salr in the wing imaginal disc might be related to the formation and/or maintenance of heterochromatic regions, at least in the wing imaginal disc. This functional aspect of *Drosophila* Sal proteins is likely conserved with their vertebrate counterparts, as human and mouse Sall1 and Sall4 proteins are preferentially bound to pericentromeric heterochromatin (Yamashita et al. 2007; Sakaki-Yumoto et al. 2006). We also found that changes in chromatin accessibility are poorly correlated to the changes in gene expression detected in *salm/salr* mutant wing discs. This observation mirrors the very limited overlap between DNA binding and mRNA expression in the case of human SALL4 (Yang et al. 2017), and the lack of correlations between changes in gene expression and chromatin accessibility in odontoblast progenitor’s mutant for *Sall1* during mouse tooth development (Lin et al. 2021).

### Salm/Salr function and heterochromatin organization

The formation and maintenance of heterochromatin in eucaryotic genomes is a fundamental aspect of the structural and functional organization of chromosomes within the nucleus. Constitutive heterochromatin is localized at subtelomeric regions and close to the centromeric DNA, is characterized by epigenetic marks (H3K9me2/3 and H4K36me2) and by global hypoacetylation of Histones, and remains mostly transcriptionally inactive (Noma et al. 2001; Grewal 2023). In contrast, facultative heterochromatin comprises genomic regions that interact with the nuclear envelope (lamina-associated domains) and nucleolus (nucleolus-associated domains), where gene silencing is regulated (Fortuny and Polo 2018). Heterochromatin formation is required for genome integrity, transcriptional silencing of repetitive DNA sequences and the maintenance of nuclear membrane stiffness (Janssen et al. 2018). Some of the alterations we found in *salm/salr* mutant cells, such as the augmented expression of *w^+^* sub-telomeric insertions, can be directly ascribed to faulty heterochromatin assembly. In addition, the appearance of mitotic defects, increased expression of retrotransposons and alterations in H3 phosphorylation observed in early blastoderms derived from *sal* heterozygous flies suggest a role for Salm/Salr in the initial establishment of centromeric heterochromatin regions. Other *Drosophila* transcription factors, such as Eyegone (Eyg) and Homothorax (Hth) have been shown to participate in the regulation of heterochromatic assembly in early *Drosophila* blastoderms (Hth; Salvany et al., 2009; Zaballos et al., 2015) or in heterochromatic gene silencing (Eyg; Salvany et al., 2012; Blom-Dahl and Azpiazu, 2018). In the case of Hth, it has been proposed that maternal Hth contributes to the transcription of satellite repeats, whereas Eyg interacts with HP1 to maintain a closed heterochromatin-like conformation (Salvany et al. 2012, 2009). The analysis of the mechanisms by which Salm/Salr affects heterochromatin formation and or maintenance are beyond the scope of this work. Thus, although the variety of genetic and developmental alterations detected in diploid and polyploid *salm/salr* mutant cells points to a role of the corresponding proteins in heterochromatin formation, the molecular events linking chromatin organization and Salm/Salr function are still unexplored.

We can envision several possible roles of Salm/Salr promoting heterochromatin formation. First, this function could be mediated by interactions with the ATP-dependent chromatin-remodeling and histone deacetylase complex (NuRD). Thus, the activities of NuRD and vertebrate Sall proteins have been linked in multitude of cellular contexts (Wang et al. 2023; Basta et al. 2017; Ma et al. 2018), and a function of NuRD in the assembly of pericentromeric heterochromatin has already been suggested in proliferating B cells and in *Schizosaccharomyces pombe* (Creamer et al. 2014; Sims and Wade 2011; Chadwick et al. 2009). Furthermore, Sall4 promotes reprogramming of mouse embryonic fibroblast by recruiting NuRD to close open chromatin (Wang et al. 2023), further suggesting a relationship between Sal function and NuRD regulating chromosomal architecture. In this context, the function of Salm/Salr would imply the recruitment to genomic targets of NuRD complexes to promote heterochromatin compaction and gene silencing. Complementary, the function of Salm/Salr might involve interactions with Lamins, which constitute the main component of the internal nuclear envelope and are also located in the nucleolus periphery. In this model, which has already been suggested for the DNA-binding protein Oct1 (Guelen et al. 2008; Malhas et al. 2009), Salm/Salr would participate in tethering heterochromatic DNA to Lamin in both the nuclear envelope and the nucleolus periphery. In fact, nuclear lamina-associated DNA domains are regions enriched in A/T sequences (Meuleman et al. 2013), the most prominent DNA binding signature of Sal proteins (Yamashita et al. 2007; Kong et al. 2021). In both scenarios; recruitment of NuRD to promote chromatin compaction and bridging chromosomal DNA to the lamina, we can expect that loss of Salm/Salr activity will lead to de-repression of gene expression, retrotransposon expression and the appearance of DNA damage (Nikolov and Taddei 2015; Chen et al. 2016). Similarly, the changes we found in the size of the nucleolus and the circularity of the nuclear envelope in salivary glands could also be caused by defects in heterochromatin formation, or in the proper attachment of heterochromatin to the nuclear envelope or nucleolus periphery (Fig. 7).

**Figure 7:**
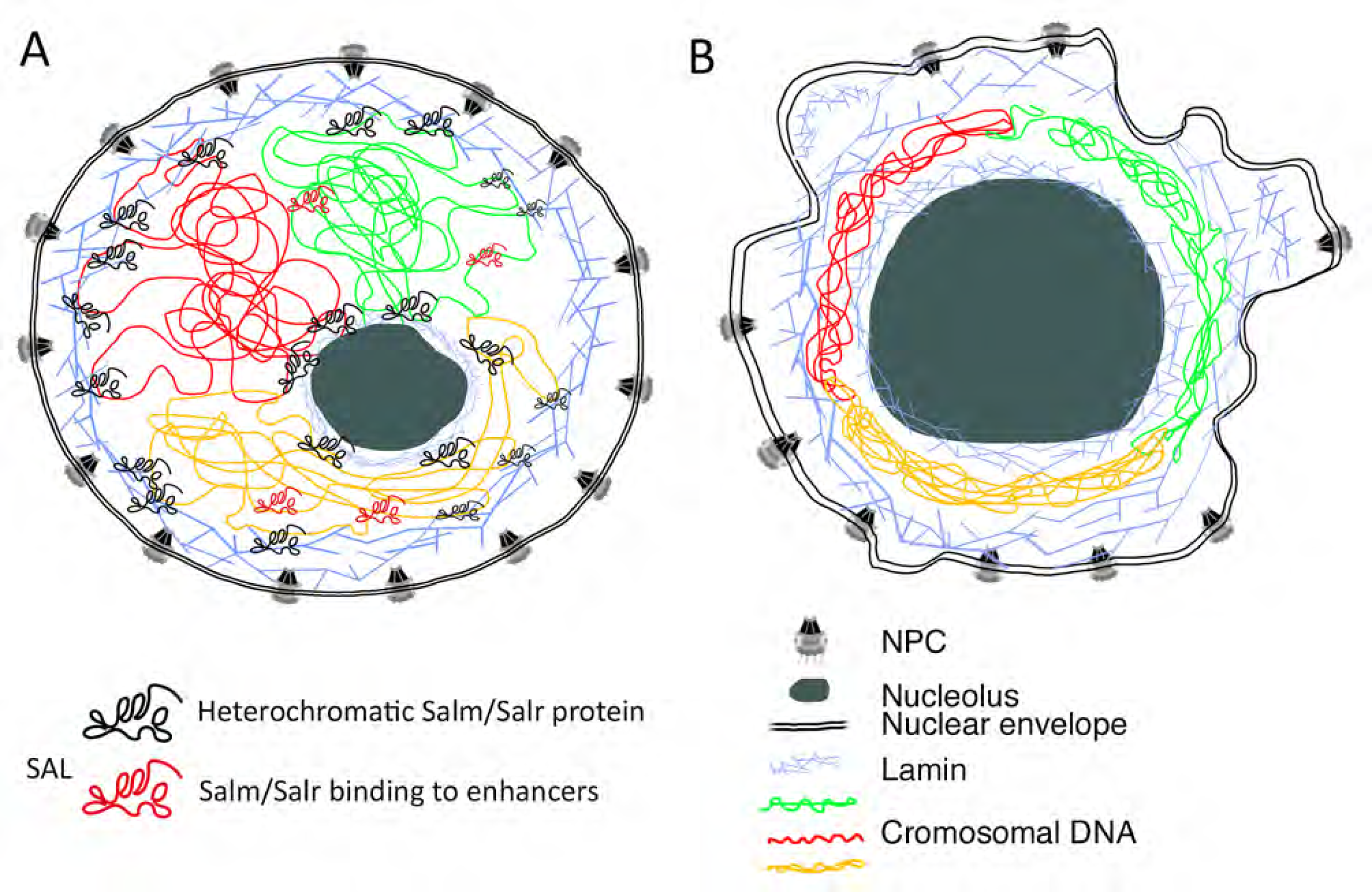
Proposed model for Salm/Salr function. (A-B) Cartoon representing wild type (A) and Salm/Salr mutant (B) nuclei. The nuclear envelope is drawn as a double line showing the nuclear pore complexes. The localization of Lamins is drawn as crisscross blue lines surrounding the nuclear envelope and the nucleolus (grey round shape). The chromosomal DNA is represented by red, blue and yellow lines within the nucleus. The Salm/Salr proteins are represented as back and red shapes, indication Salm/Salr binding to euchromatic regions and acting as a canonical sequence specific transcription factor (red) or as a heterochromatic binding protein (black). The main alterations observed in *salm/salr* mutant polyploid cells are shown in B, including wiggly nuclear envelope, enlarged nucleolus and reorganization of nuclear DNA around the periphery of the nucleus.

### A function of salm/Salr preserving genome integrity

The organization of DNA in chromatin domains has a broad impact on the maintenance of genome integrity, influencing both the susceptibility to DNA damage and the efficiency of DNA repair (Ortega et al. 2021). In fact, changes to the heterochromatin landscape can be associated to mitotic errors, aberrant chromosomal structures, replication stress and replication-transcription conflicts associated to increased transposable elements transcription (Janssen et al. 2018). It was known that *salm/salr* mutant cells display low levels of apoptosis and are preferentially stalked in the G2 phase of the cell cycle (Organista and de Celis 2013), two cellular phenotypes that we now can relate to the generation of DNA damage in these cells. Thus, we were able to visualize DNA damage in *salm/salr* mutant cells, and also detected increased expression of P53 in these cells. Consistently, there is a considerable overlap between the transcriptional response to irradiation and the changes in gene expression observed in *salm/salr* mutant wing discs. The expression of genes induced by irradiation and overexpressed in *salm/salr* mutant discs requires P53 (van Bergeijk et al. 2012), and includes several genes encoding enzymes involved in the response to DNA damage that are not normally expressed in the wing disc. We have not explored the mechanistic details leading to the activation of a p53-dependent DNA damage response in *salm/salr* mutant cells. However, considering other aspects of the *salm/salr* phenotype, we suggest that DNA damage and the corresponding response occur as a consequence of the function of Salm/Salr proteins in the maintenance of heterochromatic domains. Interestingly, several links between Sal function and DNA damage have already been identified, including the induction by Sall4 of resistance to irradiation in a P53 dependent manner (Nie et al. 2019). Certainly, the interrelationships between DNA integrity and Sal proteins requires much additional research, and could be a determining factor in the function of these genes in the development of embryonic stem cells and human cancers associated to altered expression of Sall1 and Sall4 (Xiong et al. 2015; Nie et al. 2019).

## Material and Methods

### Drosophila melanogaster cultures

Flies were kept at 17 °C or 25 °C in temperature rooms with 55% humidity. We used standard fly food containing glucose, agar, yeast, propionic acid and flour. We used the Gal4/UAS system (Brand and Perrimon, 1993) to express *salm* and *salr* RNAi, *UAS-salm-RNAi* (VDRC3029) and *UAS-salr-RNAi* (VDRC28386) in the wing disc (*sal^EPv^-Gal4*; Cruz et al., 2009) and the salivary gland (*AB1-Gal4* and *sal^EPv^-Gal4*). We also used UAS-mCD8-GFP and the 2R and 3L mini *w^+^* sub-telomeric insertions P[+mW.hs] (BL44260 and BL44261). We also used a chromosomal deficiency for the *salm* and *salr* genes (Df*(2L)32FP5*; (Barrio et al. 1999).

### Gene expression analyses

We selected a dataset of 1622 genes which expression changes comparing imaginal wing discs of *sal^EPv^-Gal4 UAS-GFP; tub-Gal80^ts^/UAS-GFP* and *sal^EPv^-Gal4 UAS-GFP/UAS-salm-i; tub-Gal80^ts^/UAS-salr-i* genotypes grown for 24 or 48 h at 29 °C before dissection in the third larval instar (Organista et al. 2015). The expression of these genes was either increased (822 genes) or decreased (700 genes) at both time points (265 and 243, respectively) or only after 24 h (60 and 105) or 48 h (597 and 352) with a logFC magnitude of −0.22 to −4.54 for upregulated genes and 0.22 to 5.26 for downregulated genes, and with p-values lower than 0.04 (Supple. Table 1). Wing imaginal disc *in situ* hybridization data in wild type and *sal^EPv^-Gal4 UAS-GFP/UAS-salm-i; UAS-salr-i*/+ genotypes were obtained from Supplementary Figures S3 to S13 included in Organista et al., 2015.

### Transposable elements expression analysis

RNA from 0-2 hours old embryos was extracted using the GE Healthcare extraction kit. 1 µg of total RNA was treated with DNAaseI at 37°C for 30 mins and retrotranscribed with the Super Script III Reverse Transcriptase (Invitrogen). The real time PCR was performed with a Bio-Rad CFX Opus 384.

The primers we used were:

-F-Element Fwd: 5‘-AGATCCGGCAGACATTCAG-3’

-F-Element Rv: 5‘-ACTTGACCATGTTTCCCCC-3’

-TART-A Fwd: 5‘-TCTCCTCAGACATGTCCCTCCCAT-3’

-TART-A Rv: 5‘-TCTTGTAGCGGCAGTTGCTAGTGT-5’

-Het-A Fwd: 5‘-TTGTCTTCTCCTCCGTCCACC-3’

-Het-A Rv: 5‘-GAGCTGAGATTTTTCTCTATGCTACTG-3’

-Rpl32 Fwd: 5‘-AGCATACAGGCCCAAGATCG-3’

-Rpl32 Rv: 3‘-TGTTGTCGATACCCTTGGGC-3’

### Quantification of eye pigment

15 heads of each genotype were frozen at −20 for 5 days and then homogenized in methanol acidified with 0,1M HCl (1:9) to extract the pigment. The extracted pigment was measured at 480nm. Three different measurements are performed for each genotype.

### ATAC-Seq

For ATAC-Seq (“Assay for Transposase-Accessible Chromatin using sequencing”) experiments, 20 imaginal wing discs were used per sample to make two technical replicates, and the larvae were of *sal^EPV^-Gal4 UAS-salm-i/+; UAS-salr-i/+* and *sal^EPV^-Gal4 UAS-GFP/+* genotypes. The discs were dissected in pre-cooled PBS and pipetted in PBT (PBS, 0.1 % Triton X-100) containing 1x proteases inhibitors (cOmplete™ *Protease Inhibitor Cocktail tablets*, Roche, 11873580001). The disks were incubated in PBT with 1x protease inhibitors and 0.4 % IGEPAL CO-630 (Sigma, 542334) for 30 m on a nutator at 4 C. The suspension was centrifugated 5 m a 4 C at 3200 g after a 30 m incubation at 4 C. The pellet was resuspended in transposition solution containing 5 μL TDE1 (Illumina, 15027865) 25 μL 2x buffer (Illumina, 15027866), 20 μL nuclease-free H_2_O and incubated 30 m at 37 C. We then added 50 μL of stop solution (50 mM Tris-HCl (pH 8.0), 100 mM NaCl, 0.1 % SDS, 100 mM EDTA (pH 8.0)) and 5 μL of RNAse at 1 mg/mL, followed by an incubation of 10 m at 55 °C. Finally, we added 3 μL of Proteinase K (20 mg/mL) and incubated 1 h at 65 °C. The DNA was purified using the DNA clean and concentrator kit (Zymo, D4014) and the columns were eluted two times in 12 μL of EB buffer. The DNA concentration was quantified with Qubit (Invitrogen). The DNA was amplified by PCR using the Nextera DNA sample preparation and indexing kits (Illumina, FC-121-1030, FC-121-1011) as follows: 20 ng of DNA, 2.5μL of i5, 2.5μL of i7 oligonucleotides, 2,5 μL Nextera cocktail and 7,5 μL of Nextera PCR mastermix (NPM, Illumina, 15027920) in a total volume of 25 μL. The PCR amplification program was 72 C 3 min, 98 °C 30 s, (98 C 10 s, 63 C 30 s, 72 C 3 min) x 12 cycles. The DNA fragments were size-selected with AMPure XP beads using 0.5 – 1.8x (Beckman Coulter, A63881) and DNA was eluted in a final volume of 15 μL. The samples were sequenced with Illumina NextSeq500, 75 bp paired-end to a depth of 50000 reads at the EMBL GeneCore.

### H3K27ac chromatin immunoprecipitation

For ChIP-seq experiments, 120 imaginal discs were used per samples. Discs were fixed in 1.8 % formaldehyde, snap frozen and store at −80 until required. *Chromatin preparation from frozen Drosophila imaginal wing discs*: the frozen disks were thawed on ice and 500 ?L RIPA buffer (140 mM NaCl, 10 mM Tris-HCl pH 8.0, 1 mM EDTA, 1% Triton X-100, 0.1% SDS, 0.1% Na-deoxycholat) and 1x Roche cOmplete Protease inhibitors. The samples were then sonicated in the Bioruptor Pico using 1.5 ml sonication tubes (Diagenode C30010016) according to the manufacturer’s instructions for 12 cycles (30 s on / 30 s off). The supernatant was transferred into 1.5 ml low binding tubes (Eppendorf, 0030108051) and centrifuged at 20.000 xg for 10 m at 4 C. For quality control and quantification of the chromatin, an aliquot of 10 μl was taken out and processed as described below. The rest of the supernatant was snap-frozen in liquid N2 and stored at −80 C until the ChIP was performed. For quality control of the chromatin, one aliquot of 10 μl was RNase treated (50 μg/ml final) at 37 C for 30 m and reverse cross-linked using a final concentration of 0.5% SDS and 0.5 mg/ml Proteinase K over night at 37 C for 10 h and 65 C for 8 h on a thermomixer with interval shaking. The next day the DNA was purified with Phenol-Chloroform purification and precipitated with ethanol, Sodium Acetate pH 5.3 and glycogen to obtain pure DNA. The size distribution was assessed by gel electrophoresis through running the DNA on a 1.2 % agarose TAE gel. The majority of the DNA was concentrated between 250 and 500 bp. The concentration was measured using Qubit hs DNA (Thermo Fisher Scientific, Q33230).

*H3K27ac* ChIP: ChIP-seq was performed as described in (Bonn et al. 2012). 1μl of H3K27ac antibody (AbCam, ab4729), was incubated overnight with the chromatin in RIPA buffer in a total volume of 900 μl. The next day 25 μl of magnetic protein A/G beads (Dynabeads, Invitrogen, 10002D and 10004D) were washed with 1ml of RIPA buffer and added to the IPs for an additional 3 h incubation on the rotating wheel at 4°C. The ChIPs were then washed for 10 min on the rotating wheel with 1x 1 ml RIPA, 4x 1 ml RIPA-500 (500 mM NaCl, 10 mM Tris-HCl pH 8.0, 1 mM EDTA, 1% Triton X-100, 0.1% SDS, 0.1% Na-deoxycholate), 1x 1 ml LiCL buffer (250 mM LiCl, 10 mM Tris-HCl pH 8.0, 1 mM EDTA, 0.5% IGEPAL CA-630 CA-630, 0.5 % Na-deoxycholate) and 2x 1 ml TE buffer (10 mM Tris pH 8.0, 1 mM EDTA) in the cold room. The chromatin was then RNase-treated and reverse cross-linked as described for the quality check of the chomatin. The molecular barcoded ChIP-seq library was prepared with all ChIPed-DNA obtained using the NEBNext UltraII DNA library Prep kit (NEB, E7645S). The quality of the libraries was assessed on a Bioanalyzer (Agilent), and libraries displayed a peak around 350-600 bp. ChIP-seq libraries were paired-end sequenced with 75 bp paired-end reads using Illumina NextSeq 500 platform at the EMBL Genomics Core Facility.

### Transmission electron microscopy (TEM)

Third instar larvae of *phm-Gal4 UAS-GFP UAS-GFP* (25 °C), *phm-Gal4 UAS-GFP Gal80^ts^ UAS-salm-i UAS-salr-i* (2 days at 25 °C, 5 days at 17 °C and 5 days at 29 °C) and *UAS-salm-i/+; AB1-Gal4*/*UAS-salr-i* (25°C) genotype were fixed in Formaldehyde: Glutaraldehyde (4%/0.04%) 2 h at room temperature and then kept at 4 °C for at least 4 days. The ring gland and the salivary gland were placed in epoxy resin, sectioned with an ultramicrotome, and the images taken in a JEM1010 TEM at 80 kV with a TemCam F416 camera and EMenu software.

### Inmunocitochemistry

Imaginal discs and salivary glands were dissected in PBS, fixed 20 minutes in PBS/TritonX100 0.3%, pre-absorbed 2 hours in PBS/TritonX100 0.3%/BSA 0.5% and incubated overnight in primary antibodies. We used rabbit anti-Salm (Barrio et al., 1996), mouse anti-Fibrilarin (Sigma), anti-H3ac, anti-HP1, anti-H3K27ac (Hybridoma bank at Iowa University), and DAPI (Merk). Secondary antibodies were from ThermoFisher Scientific (used at 1/200 dilution). Confocal images were captured using a LSM710 confocal microscope. All images were processed with the program ImageJ2 v2.3.0/1.53q (NIH, USA) and Adobe Photoshop 24.7.0.

### Comet assay for wing imaginal disc cells

A total of 40 wing imaginal discs cells per replicate were dissected in pre-cooled PBS and incubated 20 min in TrypLE^TM^ (Thermo Fisher Scientific, Waltham, Massachusetts, USA) at 37 °C. We added PBS (1:1) and centrifugated at 200 g for 10 min. The discs were resuspended and washed in PBS twice and stored at −80 °C in a Citrate buffer containing 10% DMSO until use. Approximately 10^4^ cells were embedded in low melting agarose (0.6%) and deposited on precoated slides with 1 % agarose. After agarose solidification (10 min on ice), the samples were incubated overnight at 4 °C in a lysis buffer containing 2.5 M NaCl, 100 mM EDTA, 10 mM Tris, 1 % Triton X-100, pH 10. The DNA was allowed to unwind for 40 min in alkaline buffer (300 mM NaOH, 1 mM EDTA; pH > 13) and the electrophoresis was carried at 0.73 V/cm. Slides were neutralized in PBS and stained with GelRed (Thermo Fisher Scientific). Samples were examined with an Olympus BX-61 microscope (Tokyo, Japan) equipped with an Olympus DP70 camara to capture 20-25 field images that were scored with the free CometScore 2.0 software (TriTek corp., Virginia, USA) to obtain the % of tail DNA of at least 200 nucleoids per mini-gel. The tail moment (tail intensity × length summed over the whole extent of the tail) was used to determine DNA damage.

### Bioinformatic analyses

The analysis of H3K27ac and ATAC data was carried out using the Galaxy platform and the DiffBind (Deseq) method for comparisons between controls and *salm/salr* knockdowns. The *Drosophila* reference genome was Dm6 (The Berkeley Drosophila Genome Project BDGP). Sequence alignment was carried out with Bowtie (v2.3.1). Sequence comparisons between samples and replicates was carried out using the Galaxy platform (The Galaxy Community et al., 2022). We used the databases OregAnno (Lesurf et al., 2016), RefFlat (Universidad de California, Santa Cruz, http://genome.ucsc.edu/goldenPath), and the following ModENCODE datasets: ChIP-Seq H3K9me3 (Oregon L3, Karpen, G. Dataset ID: 4952) and ChIP-Seq HP1 (Oregon L3, Karpen, G. Dataset ID: 4936).

### Correspondence between epigenetic modifications observed in salm/salr knockdown discs and candidate affected genes

In order to associate the sequences enriched in ATAC-seq and H3K27ac ChIP-seq with genes we first used the database ORegAnno (Lesurf et al., 2016), which includes 4242 regulatory regions associated to specific genes. For all those regions without ORedAnno annotations we used a criterium of proximity (van Arensbergen et asl., 2014; Zabidi and Stark, 2016). We classified our sequences as belonging to “Promoter regions”, when there is overlap with at least one transcription start site (TSS), “Intragenic regions”, when they are located within introns and located less (Proximal) or more (Distal) than 5 kb from a TSS, and “Intergenic regions”, for all sequences located between coding regions with a TSS at less than 1Kb (Proximal), between 1 and 5 Kb (Medial) and more than 5 Kb (Distal).

### Statistical analyses

Comparisons between measures were analyzed using T-Student. The p-values were grouped as indicating moderately significant difference (p-value<0.05*), significant (p-value<0.01**) and highly significant (p-value<0.001***), indicating confidence values of 95%, 99% and 99,9%, respectively. In all genomic analyses the p-values were corrected using False Discovery Rates (FDR).

## Acknowledgments

This research was supported by Secretaría de Estado de Investigación, Desarrollo e Innovación, Grant/Award Number PID2022-141894OB-C21 to JFdC. Work in the Furlong lab is partially funded by an ERC advanced grant DeCRyPT (787611), and DFG-SPP 2202 grant to EEF. We thank the Developmental Studies Hybridoma Bank at Iowa University and Bloomington Stock Center for providing the tools necessary for this work. We would also like to acknowledge Milagrós Guerra for her help with transmission electron microscopy and the support from the *Drosophila* transgenesis and confocal microscopy CBMSO scientific services. The CBMSO enjoys institutional support from the Ramón Areces Foundation. All raw data was submitted to EMBL-EBI’s ArrayExpress public repository under accession numbers: E-MTAB-xxxx (ATAC-seq data) and E-MTAB-xxxx (H3K27ac ChIP-seq data).

## Author Contributions

CMO and JFdC designed the study, C.M.O., J.F.d.C., N.A., A.-L.P., M.M., M.R.-L., A.L.-V., R.R.V. and C.G. performed the experimental work. J.F.d.C and E.E.M.F acquired the funding, J.F.d.C wrote the original draft of the manuscript and edited it, E.E.M.F reviewed it, and C.M.O., N.A., A.-L.P., M.M. and M.R.-L. critically reviewed the manuscript. All authors approved this version of the manuscript.

